# The Relationship Between Visual Motion Detection Thresholds and Visual Sensitivity to Medial/Lateral Balance Control

**DOI:** 10.1101/2025.02.11.636904

**Authors:** Stephen J. DiBianca, Hendrik Reimann, Julia Gray, Robert J. Peterka, John J. Jeka

**Affiliations:** University of Delaware, Kinesiology and Applied Physiology, Newark, DE, United States; Oregon Health & Science University, Portland, OR, United States

## Abstract

The ability to differentiate between self-motion and motion in the environment is an important factor for maintaining upright balance control. Visual motion can elicit the sensation of a fall by cueing a false position sense. While there is evidence supporting the role of central vision and fall risk, including measures of contrast sensitivity, depth perception, and size of the visual field, the relationship between motion detection thresholds and balance control remains unknown. The aim of this study is to explore the relationship between thresholds for visual motion detection (VMDTs) of the environment and measures of visual sensitivity to balance disturbances in the environment while walking. Thirty young adults (18–35 years) and thirty older adults (55–79 years) participated in a counter-balanced study where they 1) walked on a self-paced treadmill within a virtual environment that delivered frontal plane multi-sine visual disturbances at three amplitudes (6°, 10°, and 15°), and 2) performed 100 trials of a two-alternative forced choice (2AFC) task in which they discriminated between a counterclockwise (“left”) and clockwise (“right”) rotation of a visual scene under three conditions (standing, standing with optic flow, and walking). Visual sensitivity was measured using frequency response functions of the center of mass displacement relative to the screen tilt (cm/deg) while VMDTs were measured by fitting a psychometric curve to the responses of the 2AFC task. We found significant positive correlations between measures of visual sensitivity and VMDTs for 7 out of the 9 conditions in young adults, and nonsignificant positive correlations between the two measures in older adults. VMDTs were overall higher in older adults, although not significantly in the standing condition, indicating more motion in the environment is required for older adults to consciously perceive it. VMDTs also tended to increase from standing, standing with optic flow, and walking, although not significantly between the standing and standing with optic flow conditions for both populations. We interpret the positive correlations between the two measures as an indication that individuals with lower motion detection thresholds can more accurately differentiate between self-motion and motion in the environment, resulting in lower responses from visual disturbances in the environment.

## Introduction

Vision plays an important role in balance control for standing and walking by providing information about self-motion relative to the environment via optical flow (Gibson, 1958). The ability to differentiate between self-motion and motion in the environment is an important factor in maintaining upright balance control. This ability is increasingly important for older adults as vision typically degrades over time (Saftari and Kwon, 2018) and the risk of falling increases compared to young adults (Appeadu and Bordoni, 2023). The current literature comparing fall risk to vision impairments focuses on aspects of visual acuity, namely contrast sensitivity, depth perception, and size of the visual field (Saftori and Kwon, 2018). However, it is well established that optic flow can elicit an illusion of self-motion, or vection, causing an automatic response from the motor control system to adjust for the perceived position sense. This response can hinder upright balance by adjusting the body configuration to a false position sense. Currently, there is little information on how the ability to detect visual motion in the environment may relate to the nervous system’s sensitivity to vision for balance control while walking. The goal of this study was to investigate the relationship between an individual’s ability to detect motion in the environment (visual motion detection threshold) with their sensitivity to a visual disturbance in the environment (visual sensitivity) while walking.

Optic flow can produce the perception of self-motion, and thus disturb upright balance in both standing (Jeka et al., 2010; Kiemel et al., 2006; Peterka, 2002) and walking (Franz et al., 2015; Logan et al., 2010; McAndrew et al., 2010; Reimann et al., 2018b). To our knowledge, there has been only one study that has directly compared measures of visual motion detection thresholds and their relationship to the control of upright balance. Specifically, data collected during the Salisbury Eye Evaluation (SEE Project) by Freeman et al (2008) found that motion detection thresholds (VMDTs) were associated with over 3 times higher odds of failing a single leg balance stance task when adjusted for age, sex, and race compared to the other measures of vision in the model including visual acuity, contrast sensitivity, and visual field. A review on aging vision and falls by Saftari and Kwon (2018) highlights the general finding that decreased visual acuity is associated with increased risk for falls and hip fractures. However, they emphasize that visual motion perception as a contributor to fall risk has been a critical omission in the literature.

To measure visual motion perception, a visual motion detection test provides a threshold value to measure the ability of an individual to consciously detect movement in the environment. To calculate a threshold value, perceptual responses are recorded after exposing participants to optic flow with varying movement directions and amplitudes (Turano and Wang, 1992). These thresholds have typically been measured while seated (Warren et al., 1989; Gilmore et al., 1992; Freeman et al., 2006, 2008; Conlon et al., 2017), where self-motion is eliminated and balance is not an issue. We’ve shown that VMDTs can be reliably obtained while both standing and walking (DiBianca et al., 2023). VMDTS have shown to increase in older adults compared to young adults while seated (Conlon et al., 2017; Gilmore et al., 1992; Habak and Faubert, 2000; Snowden and Kavanagh, 2006; Tran et al., 1998; Warren et al., 1989), therefore, we expect to see the same trend in standing and walking.

Measuring the sensitivity of balance to a disturbance in a sensory modality (e.g., vision, vestibular) has been explored in postural control studies using a linear systems theory approach (Cenciarini and Peterka, 2006; Jeka et al. 2010; Logan et al. 2010; Goodworth and Peterka, 2012). Presenting a sensory stimulus with a known frequency and amplitude, postural responses can be measured to understand how sensitive a participant is to a sensory stimulus. Visual sensitivity (cm/deg) has been recently measured while walking using a similar approach in which the amount of center of mass (CoM) movement (cm) was measured relative to the amplitude of tilt (in degrees) in a virtual environment (DiBianca et al., 2024). The work presented here includes data from the visual sensitivity measures reported by DiBianca et al. 2024 and compares those findings with VMDTs taken from the same cohort. Results from that work showed that sensitivity to visual disturbances was higher in older adults compared to younger adults, supporting previous findings that older adults are more sensitive to visual disturbances to balance while walking (Franz et al., 2015; Osoba et al., 2019).

Vision degrades with age. Anatomical changes of the eyes occur in which the lens becomes thicker and loses elasticity which makes it more difficult to focus on near and far-sighted objects (Saftori and Kwon, 2018). Structural changes are accompanied by functional changes including a reduction in the quality of visual inputs to the central nervous system (Owsley, 2011). We expect a stronger coupling between VMDTs and sensitivity to visual disturbances in younger compared to older adults due to older subjects having degraded quality (higher noise) in their processing of visual motion information for balance control.

The main goal of the current study was to test whether VMDTs are related to balance disturbances caused by visual motion of the environment. Three conditions of VMDTs were implemented to compare with the measures of sensitivity to visual disturbances: 1) standing, 2) standing with simulated optic flow, and 3) walking. A VMDT measured while standing may be a more accurate representation of the visual system’s ability to detect motion since the head is stationary, however a VMDT measured while walking presents higher ecological validity for exploring fall risk to a visual disturbance while walking. It is currently unclear whether measures taken during standing versus walking differ in their relationship to the susceptibility of visual-evoked balance disturbances while walking. We expected thresholds to increase from standing to standing with optic flow, and then a further increase in walking due to the added noise (variability) from both optic flow motion and from head movement.

According to the sensory reweighting theory, the nervous system can be thought of as a maximum-likelihood integrator which takes a weighted combination from each sensory pathway to produce an optimal estimate that has lower variance than either of the individual sensory signals (Ernst and Banks, 2002; Peterka, 2002). Our expectation was that the ability to better detect motion in the environment (lower threshold) would be indicative of a more reliable visual system. An improved ability to detect environmental motion (lower threshold) would bias individuals towards greater reliance on visual cues for estimates of self-motion with the result that they rely more on vision for balance (higher visual sensitivity). Therefore, we expected to see an inverse correlation between individuals’ VMDT and measures of visual sensitivity.

We hypothesize that the measured VMDTs will: 1) inversely correlate with sensitivity to a visual disturbance while walking, 2) increase from standing, to standing with simulated optic flow, to walking, 3) be larger in young adults compared to older adults, and 4) will have a stronger relationship in young versus older adults.

## Research Methods and Design

Thirty healthy young participants between the ages of 18-35 as well as thirty healthy older participants between the ages of 55-79 were recruited for this experiment for a total of 60 participants. Participant demographics are highlighted in Table 1. A screening form excluded any participants who had: any head, neck, face, or lower extremity injury within 6 months prior to experiment date, any medication that could affect balance, any neurological or cardiovascular conditions, pregnancy, BMI over 30. All participants had normal or corrected to normal vision. Verbal and written consent was obtained from each participant. The experiment was approved by the University of Delaware Institutional Review Board.

**Table 1:**
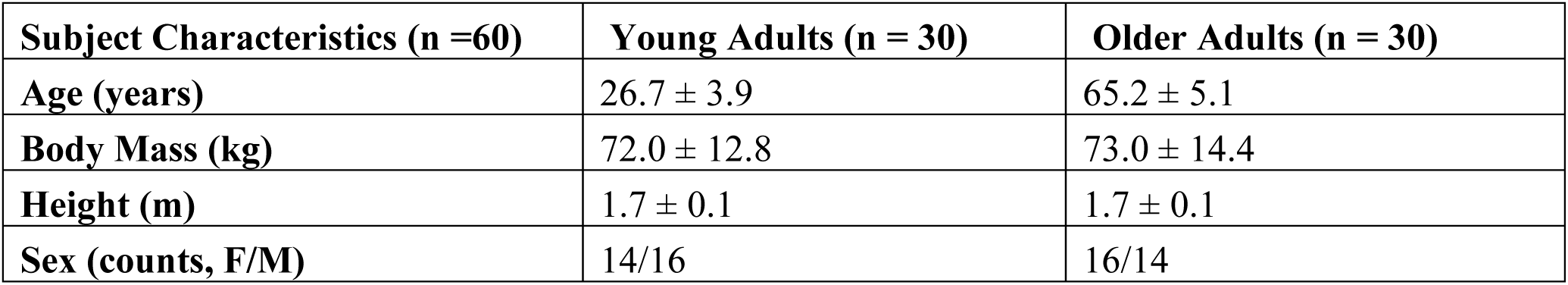
Subject Characteristics Data are presented as mean ± SD unless otherwise stated.

### Experimental Protocol & Setup

Participants performed a Snellen chart at arrival for visual acuity measures. Each participant observed a Snellen chart for each eye independently, and both eyes together at the 20/25 line and 20/20 line in which the number of letters correctly identified were summed for a maximum score of 45. One older participant did not complete a Snellen chart assessment due to experimenter error.

Participants then stood and walked on a self-paced, tied-belt treadmill (Bertec, Inc, Columbus, OH) inside a virtual environment displayed on a large dome that occupied the subjects’ full visual field. Figure 1 highlights the experimental design. All participants began with a 15-minute walking block to familiarize themselves with the virtual environment, the self-paced treadmill, the two-alternative forced choice task used to measure perceptual thresholds, and viewing the visual multi-sine stimulus used to perturb balance and measure visual sensitivity. In a counter-balanced design, participants were then selected to perform visual sensitivity testing first or perceptual testing first.

**Figure 1:**
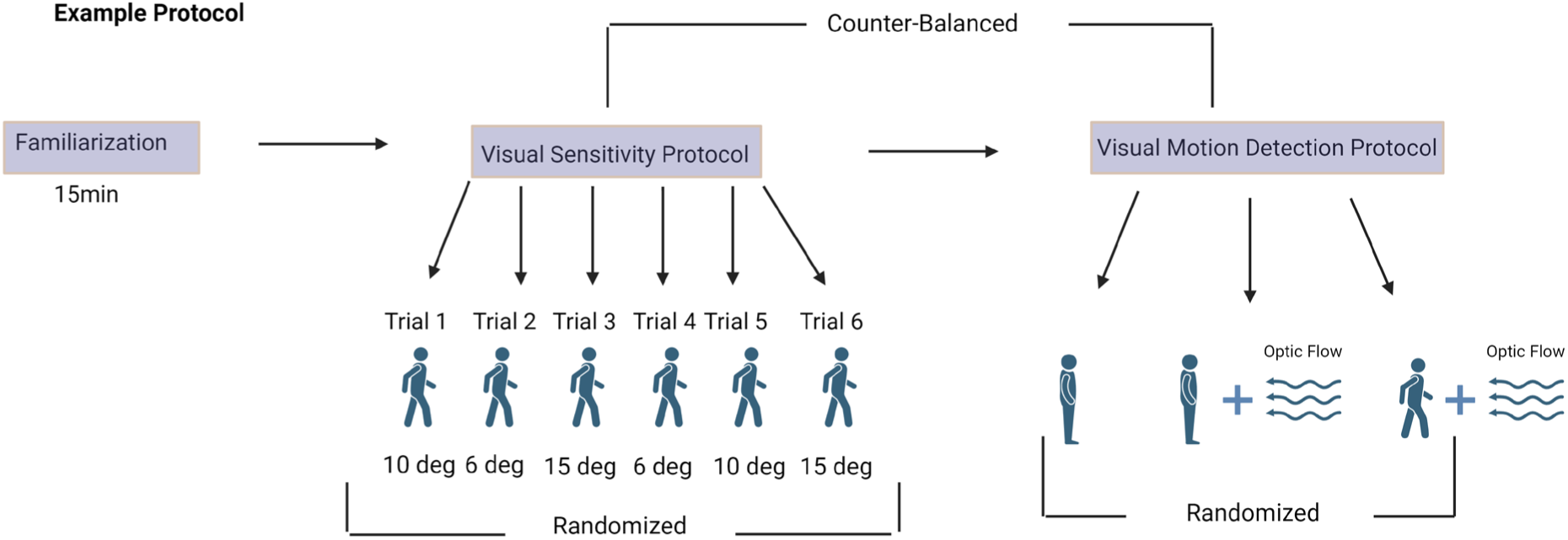
The experimental design. Participants first experienced a familiarization period where they: 1) walked on the self-paced treadmill in the virtual environment and were exposed to the visual multi-sine stimulus; 2) performed 10 trials of the two-alternative forced choice task while walking. After familiarization, participants started with the visual sensitivity protocol, or the visual motion detection protocol, which were counter-balanced. The trial order within both protocols was randomized. Created with BioRender.com

### Visual Sensitivity Measurement Protocol

Measurements of visual sensitivity were performed in a virtual walking environment (Figure 2) designed and implemented in Unity3d (Unity Technologies, San Francisco, CA, USA.). The virtual environment consisted of a tiled floor with floating cubes randomly distributed in a volume 0–10 m above the floor, 2–17 m to each side from the midline, and infinitely into the distance, forming a 4m wide corridor for the subjects to walk through. A fog was displayed in the distance to obscure the limits of the tunnel and create a perception of infinite walking distance. The anterior/posterior movement of the virtual scene matched the speed of the treadmill, and perspective in the virtual world was linked to the midpoint between two reflective markers on the subject’s temples. This perspective was linked to the center of the horizon and was the axis of rotation for both the visual multi-sine stimulus.

**Figure 2:**
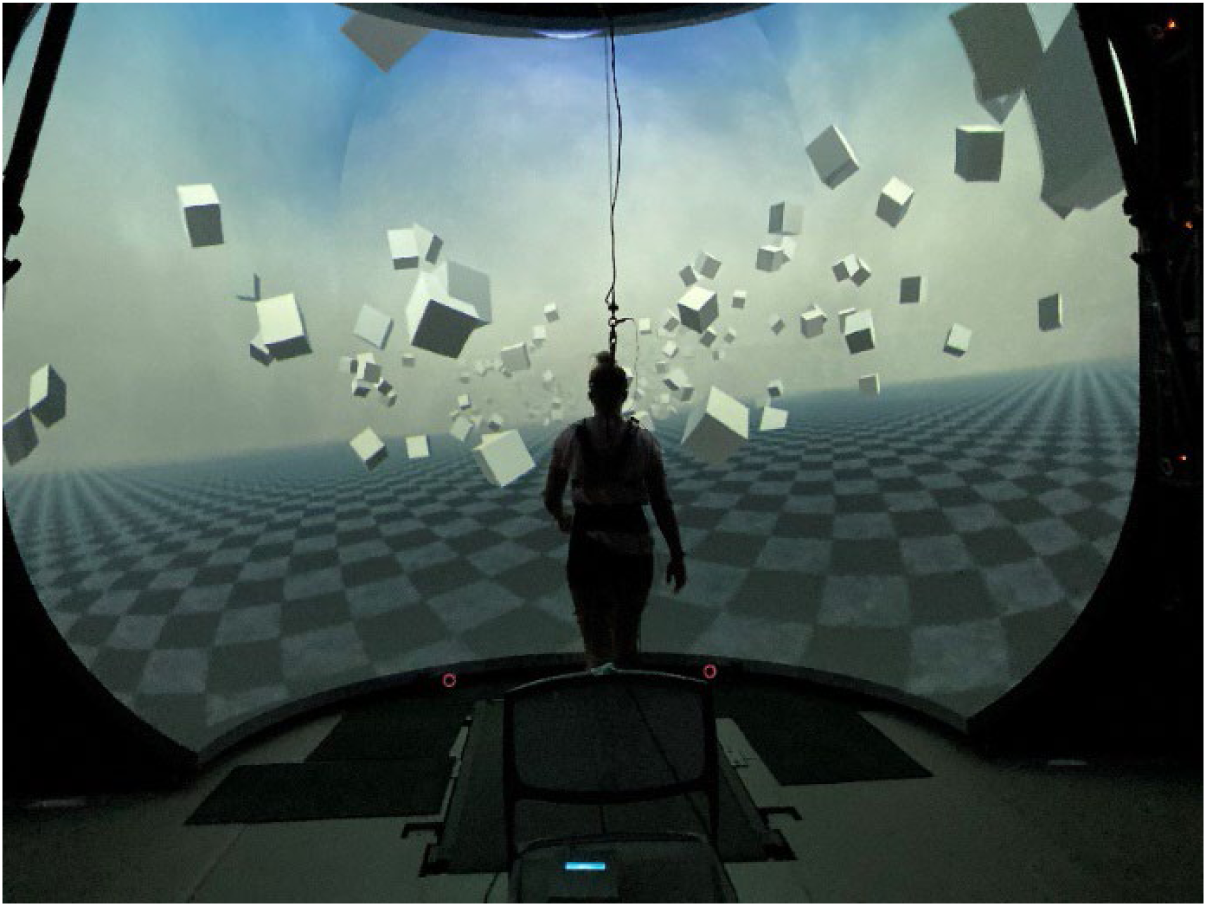
The experimental set up for the visual sensitivity measure. Healthy young and older adults walked on a self-paced treadmill in the virtual dome displaying the visual multi-sine stimulus which rotated the visual scene about an anterior-posterior axis at floor level that evoked medial/lateral body displacements.

Additionally, the treadmill control included a nonlinear proportional-integral-derivative (PD) controller and was implemented via Labview (National instruments Inc., Austin, TX, USA) to keep the midpoint between the two reflective markers placed on the posterior iliac spines at the midline of the treadmill. With a PD controller, the treadmill sped up and slowed down with the speed of the subject to maintain the subject in the center of the treadmill. Force plates integrated in the treadmill measured ground reaction forces and moments at a sampling rate of 1,000 Hz from both belts. Subjects wore a safety harness in the event of a fall, although none occurred.

To calculate visual sensitivity, participants walked while following a metronome at 0.8 Hz with two beats per cycle, or 96 bpm, while viewing a visual multi-sine stimulus that perturbed balance in the medial/lateral (M/L) direction. The stimulus consisted of a combination of ten sinusoids (Van der Ouderaa et al., 1988) at frequencies between 0.025 and 0.475 Hz, containing power at a wide range of frequencies typically observed in upright balance control (Yamamoto et al., 2015). The individual sinusoids were phase-shifted, scaled, and superposed to create a smooth, pseudorandom stimulus that is unpredictable for the participant and appears as a continuous seesaw-like motion of the virtual environment around the anterior/posterior axis at floor level. Each trial was 290s including 17s of unperturbed walking before stimulus onset followed by 6 cycles of the visual multi-sine stimulus (40s cycle duration). Participants performed two trials of the visual multi-sine stimulus at 3 different peak-to-peak amplitudes: 6, 10, and 15 degrees. The 6 trials were presented in a randomized order.

The M/L displacement of CoM was considered the sway response output for measurement of visual sensitivity. CoM position data was calculated based on a geometric model with 15 segments (pelvis, torso, head, thighs, lower legs, feet, upper arms, forearms, hands) and 38 degrees of freedom (DoF) in OpenSim (Delp et al., 2007; Seth et al., 2018) using an existing base model (Delp et al., 1990). We used the Plug-in Gait marker set by Davis et al. (1991), forty-four reflective markers were placed bilaterally on the subjects’ feet, shank, upper legs, hips, torso, upper arms, forearms and hands, with six additional markers placed on the anterior thigh, anterior shank, and 5^th^ metatarsal. Marker coordinate positions were recorded using a Qualisys Motion Tracker System with 13 cameras at a sampling rate of 200Hz.

A Fourier transform was performed on both the visual stimulus and the average CoM response time trajectories to convert the signals into the frequency domain. A frequency response function (FRF) was calculated by dividing the cycle-averaged Fourier transform of the M/L CoM response by that of the stimulus. The FRF is a complex valued function that characterizes the dynamic behavior of a system by showing the response sensitivity (gain) and timing (phase) of the system as a function of stimulus frequency. The FRF gain function (with units of cm/deg) was calculated as the absolute value of the FRF at the 10 multi-sine stimulus frequencies. These gain values were averaged to define a measure of the sensitivity of balance to visual motion:

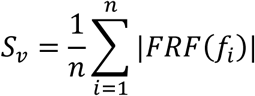

with n =10 being the number of frequencies, *f*_*i*_, included in the visual stimulus.

### Visual Motion Detection Threshold Measurement Protocol

Visual motion detection thresholds were measured under three different conditions (Figure 3): standing, standing with simulated optic flow, and walking with optic flow. The visual scenes consisted of 1,000 cubes floating before a dark background, randomly distributed in a cylindrical tunnel along the anterior-posterior axis with a radius of 14-40 m from the central axis through the treadmill. Each cube was 1×1×1 m in size. A red sphere was linked to the midpoint between the two markers placed on the temples as a focal point for participants and was placed 50 m ahead in the virtual environment. Fog was displayed in the distance to obfuscate the end of the tunnel and create the perception of infinite distance.

**Figure 3:**
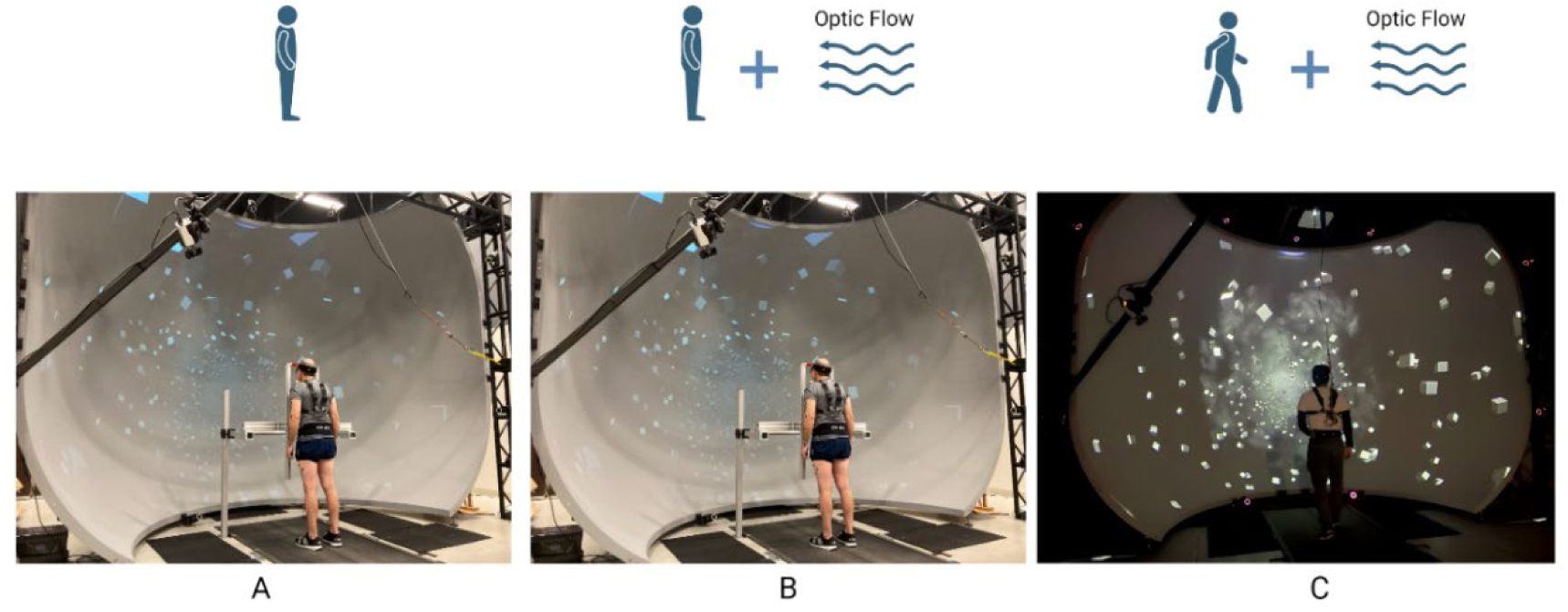
The visual motion detection experimental protocol: A) standing; B) standing with optic flow; C) walking with optic flow. For the standing conditions, participants stood with their chin resting on a head mount. Room lights were turned on in Panel A and B to better depict the chin rest although lights were off during the testing.

In the standing condition, there was no anterior/posterior optic flow. In the standing with optic flow condition, anterior/posterior optic flow was based on the baseline waking speed participants walked during the familiarization period. In the walking condition, the anterior-posterior optic flow matched the speed of the treadmill. For the standing conditions (Figures 3A&B), a custom built, adjustable head mount was used to rest their chin on the mount (without supporting weight onto it) to stabilize their head. Laboratory lights were off in all conditions.

To measure visual motion detection thresholds (VMDTs) in each condition (standing, standing with optic flow, walking with optic flow), participants performed a psychophysical test consisting of 100 trials of a two-alternative forced choice (2AFC) task (DiBianca et al., 2023). With the subject viewing the red sphere in the distance, on each trial the screen rotated either counter-clockwise, “left”, or clockwise, “right”. The rotation consisted of a single cycle of a 1 Hz raised cosine waveform with its total angular displacement defining the stimulus amplitude during which a monotone sound was played. When the sound stopped, the subject was required to verbally reported their perceived direction of motion as “left” or “right”. The stimulus amplitude was adjusted using an adaptive 3 down 1 up staircase algorithm (Karmali et al., 2016) for parameter estimation by sequential testing (Taylor and Creelman, 1967). The amplitude decreased after three correct responses and increased after one incorrect response, until 100 trials were completed.

For the standing conditions, the stimulus was manually triggered by the experimenter at an arbitrary time every one to two seconds after each response. For the walking condition, after each verbal response the experimenter triggered a custom LABVIEW program which prompted a stimulus on the third step of the right foot by tracking the peak anterior displacement of the heel marker for heel strike identification. After every 25 trials the subject was given a brief break from stimulus presentations, with each break typically lasting about 15 seconds. In the walking trials, subjects kept walking normally during these breaks. After each block of 100 trials, subjects took longer breaks of at least two minutes or more if needed, before proceeding to the next condition.

To obtain a VMDT, we fit a psychometric curve to the 100 binary responses of the 2AFC task in each condition. The fit was performed in MATLAB’s (2020A Mathworks Inc.) *fitglm* function using a generalized linear model (GLM) with a probit link. The motion detection threshold was defined as the value corresponding to a target probability of ^3^√0.5 = 0.79 (Hall, 1981; Karmali et al., 2016; Levitt, 1971; Taylor and Creelman, 1967), meaning the threshold corresponds to the amplitude of rotation at which the subject responded “right” with 79% accuracy.

Data from one older adult’s standing condition was removed as the model did not produce a reliable fit causing the threshold data point to be an extreme outlier with ∼7 times higher value than the population average.

### Statistical Analysis

To evaluate whether visual motion detection thresholds relate to visual sensitivity for balance control, and how those relationships may change due to age, we ran Spearman rank correlations between the measures of visual thresholds and visual sensitivity for both groups. Friedman tests were performed to test for any differences in visual motion detection thresholds between conditions (standing, standing with optic flow, walking with optic flow) and between groups (older and younger adults). Effect sizes were calculated using Kendall’s W. Wilcoxon tests were run to compare between groups for each condition.

## Results

All 60 participants completed the study. Average optic flow speed during the standing with optic flow condition was similar between healthy young adults (1.03 m/s ± 0.13 m/s) and older adults (1.01 m/s ± 0.14 m/s). Average optic flow speed (walking speed) during the walking condition was also similar between young adults (0.81 m/s ± 0.23 m/s) and older adults (0.81 m/s ± 0.24 m/s). Optic flow speed during the standing with optic flow condition was determined during baseline walking in the familiarization period. We see that participants walked on average slower when walking and performing the 2AFC compared to baseline walking during the familiarization period of the protocol.

### Visual Motion Detection Thresholds

Figure 4 highlights the results of the VMDTs for young and older adults for the three conditions of standing, standing with optic flow, and walking with optic flow. Young adults showed an average VMDT of 0.45 deg ± 0.33 deg, 0.45 deg ±0.29 deg, and 0.71 deg ± 0.43 deg for the three conditions, respectively. Older adults showed higher average VMDTs of 0.70 deg ± 0.65 deg, 0.77 deg ± 0.58 deg, and 1.18 deg ± 0.76 deg for the three conditions, respectively.

**Figure 4:**
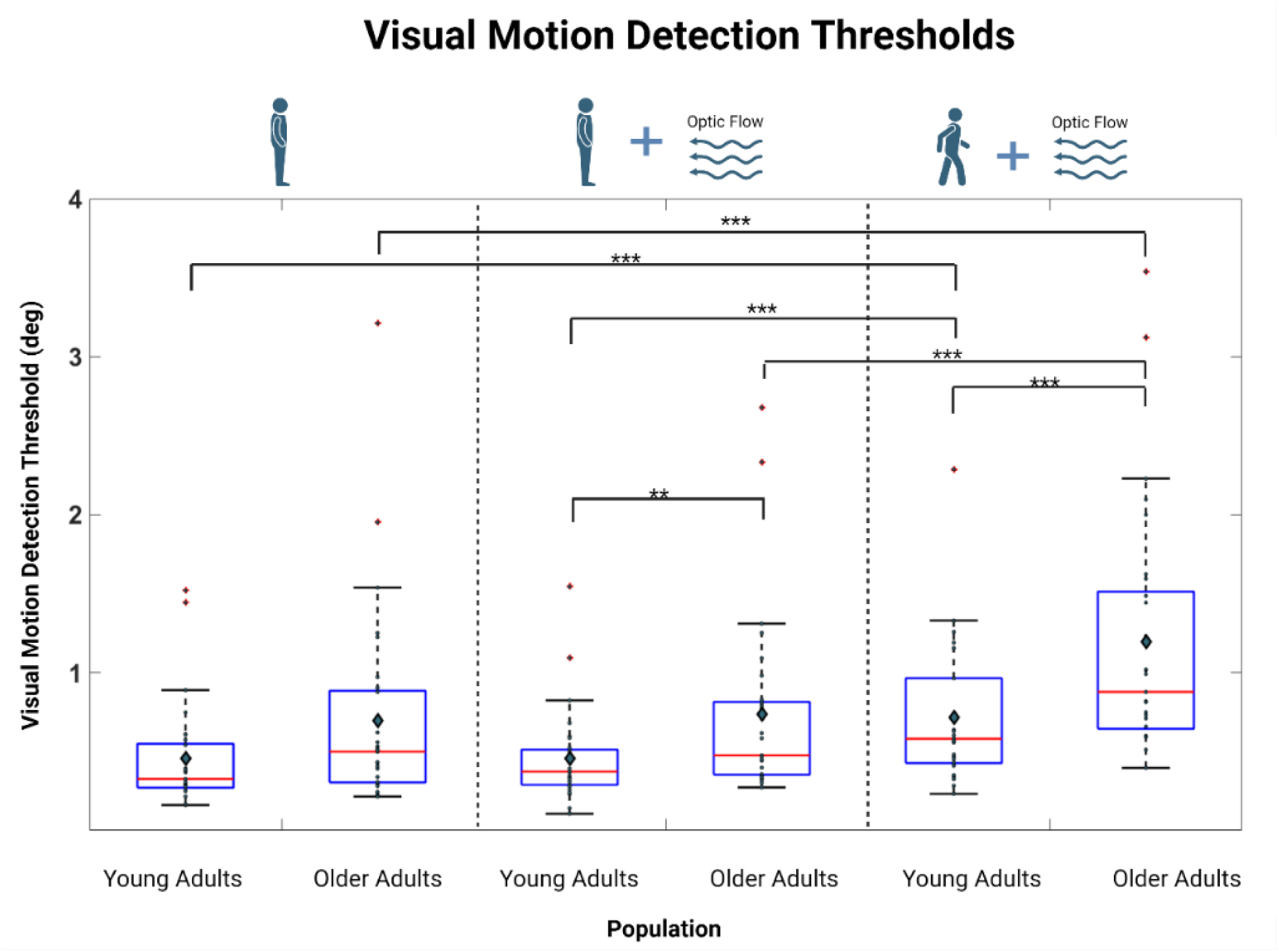
Box and whisker plots of visual motion detection thresholds measured for young and older adults while standing, standing with optic flow, and walking. The diamonds represent group means for that condition. Two stars indicate a p-value less than 0.01 and three stars indicate a p-value less than 0.001.

Results of the Friedman test for healthy young adults revealed a significant effect of condition on the resulting VMDT (χ^2^(2) = 22.2, p < 0.001) with a moderate effect size detected (W = 0.368). A follow up post-hoc analysis using pairwise Wilcoxon signed-rank tests with a Bonferroni correction revealed 1) no significant difference between the standing and standing with simulated optic flow condition (p = 0.58), 2) a significant difference between the standing and walking condition (p < 0.001), and 3) a significant difference between the standing with simulated optic flow and walking with optic flow condition (p < 0.001).

Results of the Friedman test for healthy older adults revealed a significant effect of condition on the resulting VMDT (χ^2^(2) = 28.8, p < 0.001) with a moderate effect size detected (W = 0.496). A follow up post-hoc analysis using pairwise Wilcoxon signed-rank tests with a Bonferroni correction revealed 1) no significant difference between the standing and standing with optic flow condition (p = 0.11), 2) a significant difference between the standing and walking condition (p < 0.001), and 3) a significant difference between the standing with optic flow and walking condition (p < 0.001).

Results of the Wilcoxon tests revealed: 1) no significant difference between VMDTs in young versus older participants for the standing condition (p = 0.07), 2) a significant difference between VMDTs in young versus older participants for the standing with optic flow condition (p = 0.006), and 3) a significant difference between VMDTs in young versus older participants for the walking condition (p < 0.001).

### Visual Sensitivity

Figure 5 shows box and whisker plots of visual sensitivity measures from the balance responses to M/L visual disturbances for both young and older adults. Young adults showed an average gain response of 2.24 cm/deg ± 0.93 cm/deg, 1.91 cm/deg ± 0.78 cm/deg, and 1.69 cm/deg ± 0.8 cm/deg for the 6, 10, and 15 degree peak-to-peak stimuli, respectively. Older adults showed an average gain response of 3.20 cm/deg ± 0.73 cm/deg, 2.87 cm/deg ± 0.75 cm/deg, and 2.52 cm/deg ± 0.64 cm/deg for the 6, 10, and 15 degree peak-to-peak stimuli respectively.

**Figure 5:**
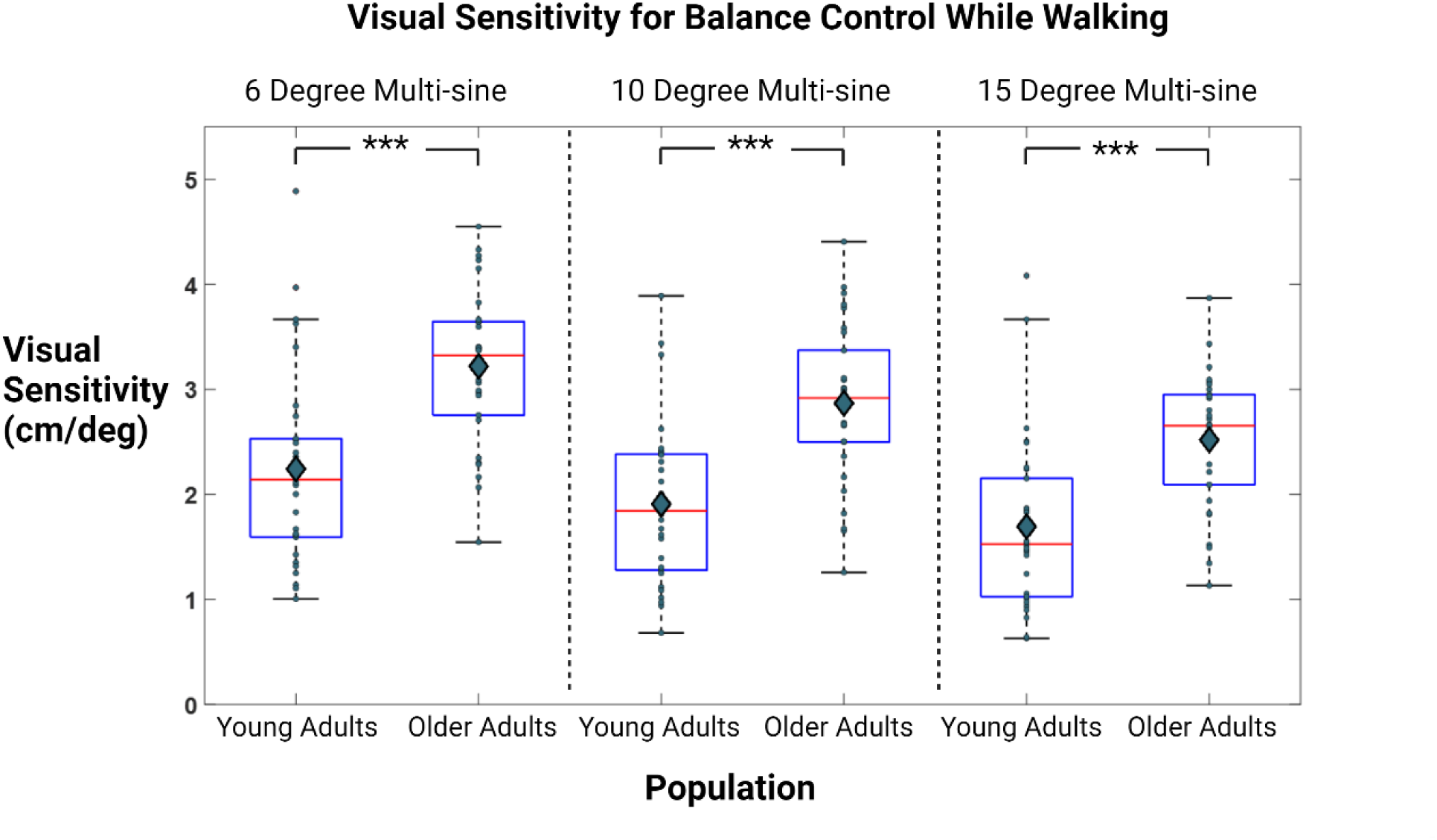
Box and whisker plots for the visual sensitivity to balance measured for young and older adults while walking. The diamonds represent group means. The *** symbols represent p values less than 0.001.

### Comparison of Visual Sensitivity and Visual Motion Detection Thresholds

Figures 6 and 7 show scatter plots relating visual sensitivity to VMDTs. All Spearman rank correlation coefficients were positive for both younger and older adults in all conditions (standing, standing with optic flow, and walking with optic flow) and for all visual stimulus amplitudes used to measure visual sensitivity (Table 2). For younger adults, the correlations were statistically significant for the standing with optic flow and the walking with optic flow conditions at all visual stimulus amplitudes, but in the standing condition the correlation was significant only at the 6 degree visual stimulus condition. In contrast, for older adults none of the correlations were significant in any condition and at any of the stimulus amplitudes used to measure visual sensitivity.

**Figure 6:**
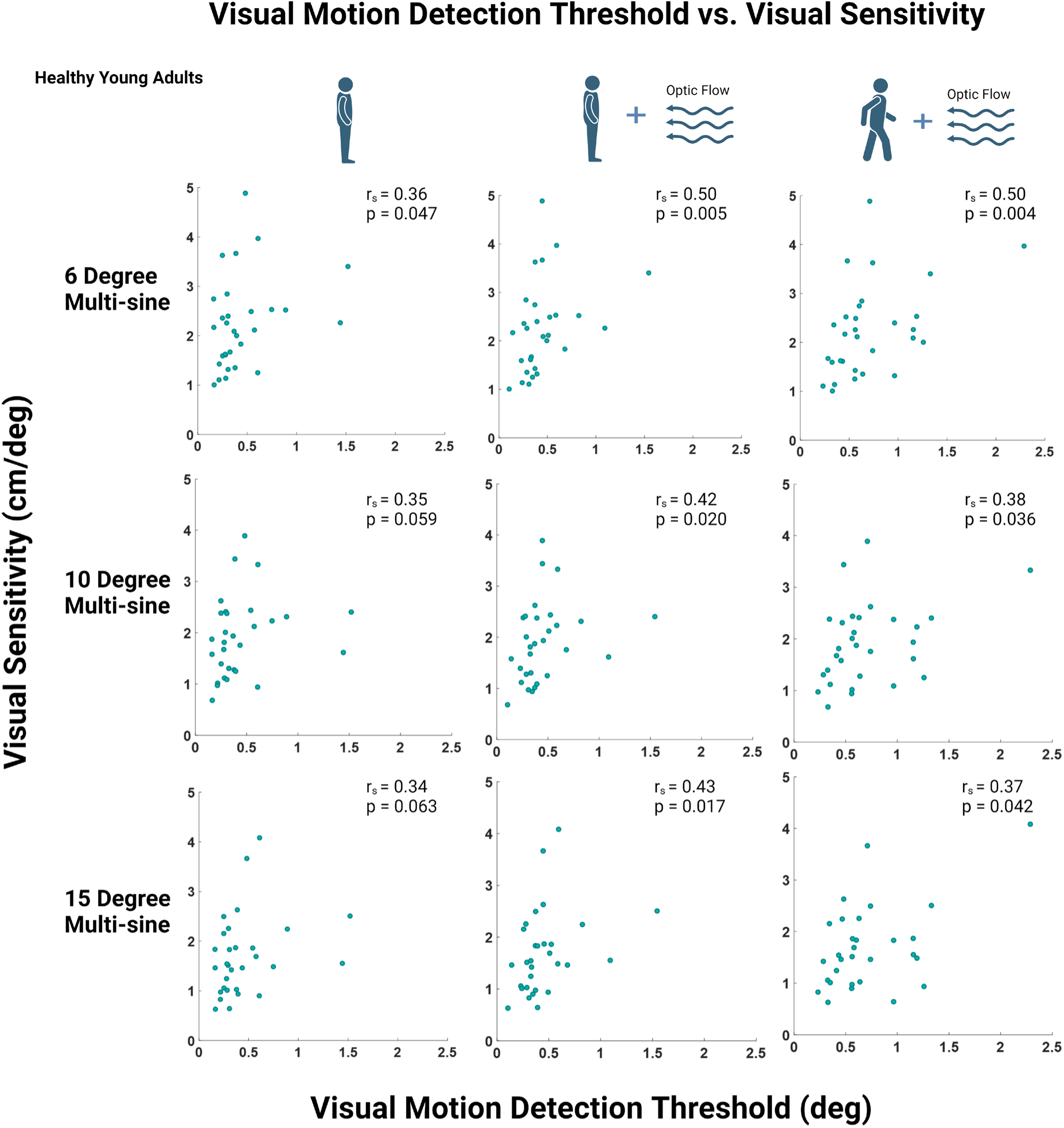
Scatter plots with visual motion detection thresholds (deg) on the x-axis and visual sensitivity to balance while walking on the y-axis (cm/deg) for healthy young adults. The 3 rows represent visual sensitivity measured during tests with 6, 10, and 15 degree stimulus amplitudes, respectively. The 3 columns represent the three VMDT conditions of standing, standing with optic flow, and walking with optic flow, respectively. Corresponding Spearman’s rank correlation coefficients (rs) and p-values are displayed above each graph.

**Figure 7:**
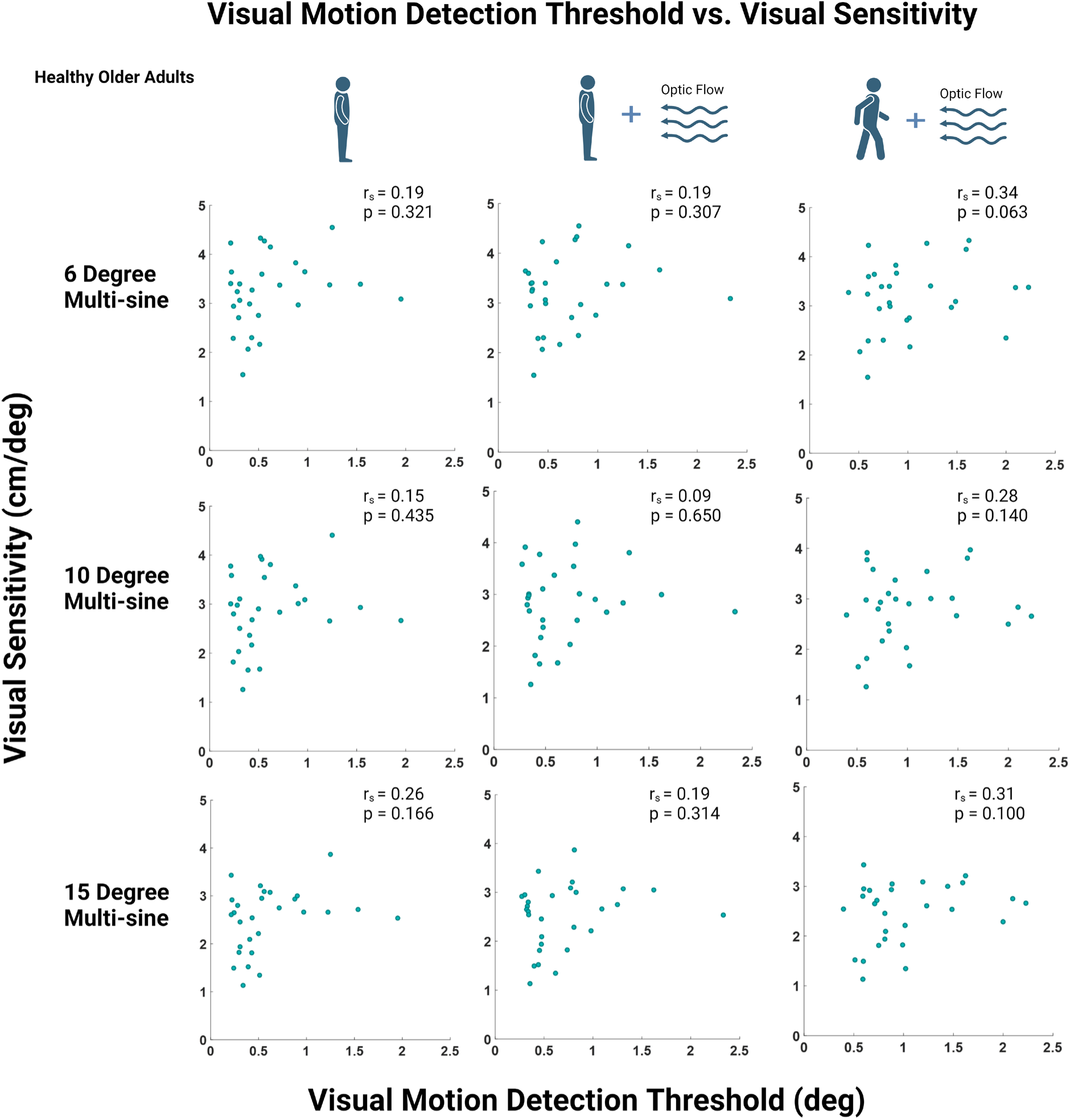
Scatter plots with visual motion detection thresholds (deg) on the x-axis and visual sensitivity to balance while walking on the y-axis (cm/deg) for healthy older adults. The 3 rows represent visual sensitivity measured during tests with 6, 10, and 15 degree stimulus amplitudes, respectively. The 3 columns represent the three VMDT conditions of standing, standing with optic flow, and walking with optic flow, respectively. Corresponding Spearman’s rank correlation coefficients (rs) and p-values are displayed above each graph.

**Table 2:**
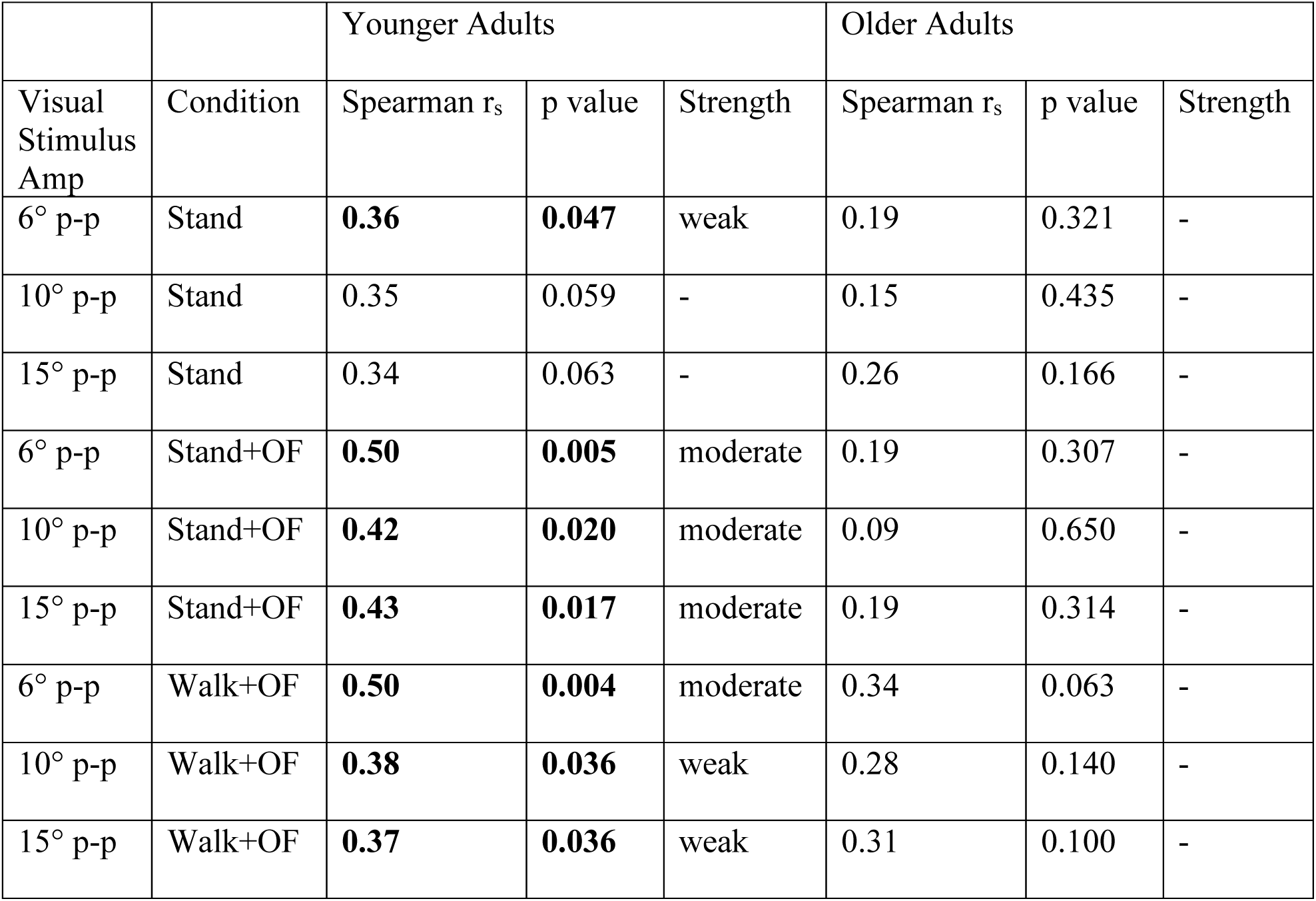
Correlations between Visual Sensitivity and Visual Motion Detection Thresholds. Qualitative ratings of correlation strengths were based on guidelines from Prion and Haerling (2014). Spearman correlation coefficients, rs, and their p-values are bolded for p<0.05. OF: optic flow, p-p: peak-to-peak amplitude of the visual stimulus.

## Discussion

Our study investigated the relationship between a participants’ sensitivity to a visual stimulus for M/L balance control and their ability to consciously detect visual motion in a virtual visual environment. Participants walked on a self-paced treadmill in a virtual environment that tilted pseudorandomly. Participants performed a two-alternative forced choice task while standing, standing with optic flow, and walking with optic flow. Psychometric fits to the experimental data estimated visual motion detection thresholds (VMDTs) for each condition. We also measured visual sensitivity to the visual motion disturbances by averaging the relative responsiveness to the visual stimulus (ratio of M/L CoM movement (cm) to the visual stimulus amplitude (deg) at the 10 driving frequencies of the visual stimulus). We found positive correlations for each of the VMDTs measured during the three conditions with the measures of visual sensitivity to balance while walking for both young and older adults. These relationships, established through Spearman rank correlations, were significant for young adults (7 out of 9 correlations) but not significant for older adults.

Our initial hypothesis was that VMDTs would be inversely correlated with visual sensitivity to balance. Our rationale was that a lower VMDT would indicate that an individual who could better detect motion in the environment would be more reliant on visual cues for upright balance control. The results showed the opposite trend and may indicate that individuals with lower VMDTs are better separating visual motion in the environment from self-motion. A lower threshold may indicate a less noisy position and velocity estimate such that those individuals are less affected by the perturbing environment because they recognize the visual motion cues are due to environmental motion and not self-motion.

Our study was motivated to investigate the influence of visual motion cues on balance. When studies investigate the relationship between visual processing and fall risk, they typically assess qualities of visual acuity. Visual acuity relates to central vision, or focal vision, capable of high spatial resolution and particularly useful for pattern and object recognition (Larson and Loschky, 2009), including measures such as contrast sensitivity (Lord and Fitzpatrick, 2001; Wood et al., 2011), depth perception (Felson et al., 1989; Lord and Dayhew, 2001), or size of the visual field (Broman et al., 2004). Visual acuity is meaningful for maneuvering around an environment and avoiding falls caused by tripping or hitting obstacles (Broman et al., 2004) as vision provides information about object size, location, and where to place the swing leg foot into a safe space. However, we know that optic flow directly influences behavioral responses due to a phenomenon called “vection,” or the visual induced perception of self-motion. Compared to central vision, the primary visual functions of the peripheral visual field include stimulus detection, flicker and motion sensitivity <u>(Monaco et al., 2007)</u>. Although not directly prompted, numerous participants reported a “zone out” strategy during the VMDT testing in which they felt better equipped to see the motion when viewing the global environment instead of focusing intently on the red sphere located in the middle of the environment. We instructed subjects to use the red dot as a guide as opposed to focusing their central attention somewhere on the outskirts of the scene. Evidence of the phenomenon of peripheral vision playing a stronger role in motion sensitivity has been supported by Haibach et al. (2009). They introduced both young and older adults to oscillating forward/backward optic flow stimuli in conditions where either central or peripheral vision were occluded and found that when central vision was occluded, vection and sway increased.

As an exploratory analysis, we ran Spearman’s correlations for Snellen chart measures of visual acuity for each participant in relation to their measures of visual sensitivity for balance control to compare the relationships between motion detection and visual acuity for visual function related to balance. These results (Supplementary Figure 1) showed no correlations between visual acuity and measures of visual sensitivity to balance while walking. There was a considerable ceiling effect with the Snellen chart measures that is particularly prevalent in young participants, as expected from a healthy population that reported normal/corrected to normal vision. Even with reported healthy vision, these results provide evidence that measures of visual function related to optic flow are a more reliable indicator for visual function related to balance control than that of visual acuity, specifically for younger adults.

Our hypothesis that younger adults would show a stronger relationship between VMDTs and visual sensitivity to walking balance control was supported. We see significant correlations in 7 out of 9 testing conditions in healthy young adults and no significant correlations in older adults (Figures 6 and 7). We interpret that this higher coupling is due to younger adults having less noise in visual inputs related to balance, providing a more salient coupling of motion processing to visual sensitivity. However, it was not expected that this relationship would be absent in older adults.

While our overall results provide a promising step in investigating the roll of visual function related to balance and potentially to fall risk, it is possible that a better measure of vection could provide a more robust biomarker into visual function related to balance control. Kooijman et al., (2023) reviewed the current literature on measures of vection which have utilized similar psychometric methods, although for the sensation of movement rather than detection of motion. The review highlights the need for an objective measure of vection. Some early promising results have been seen in electroencephalography (Keshavarz et al., 2015; McAssey et al., 2020) and center of pressure postural measures (Weech et al., 2020). Anecdotally, during threshold measurements while walking, some participants mentioned they felt themselves move even if they did not recognize the environmental motion. This self-motion recognition could have helped them identify the correct direction of the visual motion even if they did not visually perceive it. It is unknown if this self-motion sensation was experienced during the standing trials. Measuring the difference between the lowest perceived motion detected and the lowest amount of motion that elicits a postural response could provide a more robust measure of vection and may provide stronger evidence for the relationship between motion processing and visual sensitivity to balance.

We hypothesized that VMDTs would increase from the standing, standing with optic flow, and walking conditions, and would be larger in older adults compared to young adults. We found that VMDTs did not significantly change from standing to standing with simulated optic flow in both populations (Figure 4). We expected that the addition of optic flow in the standing condition could add noise to the visual system and significantly increase thresholds compared to standing without optic flow, but this was not the case. We also expected thresholds to increase from standing to walking based upon our previous results from DiBianca et al. (2023). This was supported in both younger and older groups (Figure 4). We also found that VMDTs were higher in older adults compared to young adults, although not significantly higher in the standing condition. This finding supports previous work on the change of motion processing due to aging for seated tests where balance is not required (Gilmore et al., 1992; Conlon et al., 2017).

## Conclusion

Our study found that measuring the ability of the visual system to detect movement in the environment was positively correlated with visual sensitivity to a visual perturbation for balance control. This correlation was significant for young adults (7 out of 9 conditions; Figure 6) but not for older adults (Figure 7). Our interpretation for this finding is that individuals with lower motion detection thresholds can more accurately differentiate between self-motion and motion in the environment, resulting in lower responses from visual disturbances in the environment. Evidence for this is supported by the fact that older adults displayed higher VMDTs and were more sensitive to visual disturbances. A lack of significant correlations in the relationship between VMDTs and visual sensitivity to balance in older adults merits the need of a more robust biomarker for vection as highlighted in a recent review by Kooijman et al., (2023). An exploratory analysis revealed that measures of visual motion detection were more robust in predicting whether an individual’s balance would be sensitive to visual disturbance in the environment than measures of visual acuity via Snellen Chart measures.

## Supplementary Material

## Supplementary Material

**Supplementary Figure 1:**
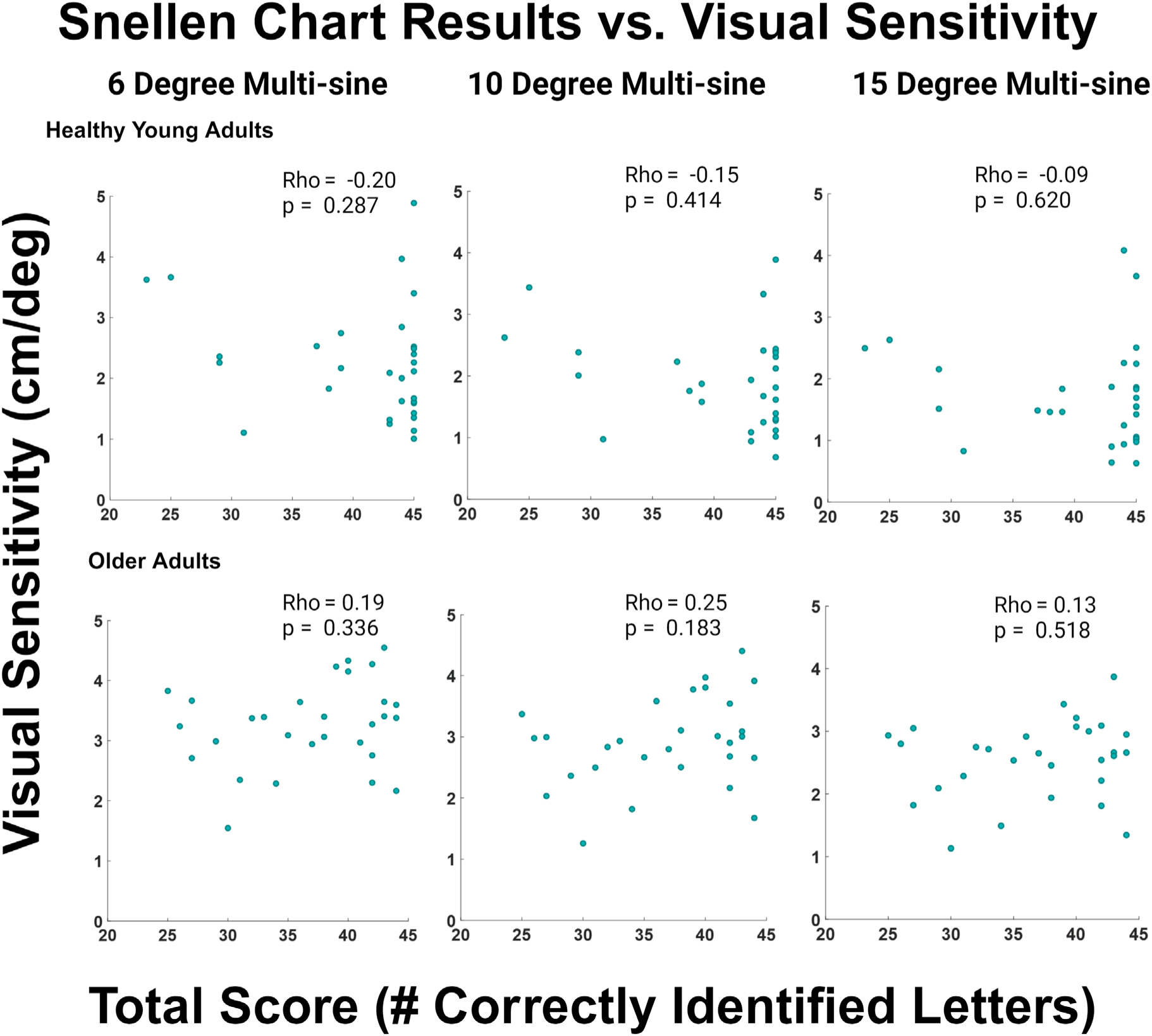
Scatter plots of Snellen chart results compared with measures of visual sensitivity to balance for both young and older adults.

